# Dye-free retinal angiography using blood-oxygenation modulation

**DOI:** 10.1101/485839

**Authors:** L.E. MacKenzie, T.R. Choudhary, J. Fernandez Ramos, N. Benjamin, C. Delles, A.R. Harvey

## Abstract

Fluorescence angiography (FA) is widely used for studying and diagnosing abnormalities in the retinal blood circulation, but has associated risks of nausea, skin irritation, and even death. We describe a new non-invasive angiography technique: Blood Oxygenation Modulation Angiography, in which multispectral imaging of a transient perturbation in blood-oxygen saturation, yields angiography sequences similar to FA, including key features such as sequential filling of choroidal and retinal-vessels, which underpin assessment of circulation health. This is the first non-invasive angiography technique capable of visualizing these circulation features.

## 1. Introduction

Fluorescence angiography (FA)[1] is widely used to study retinal blood circulation, enabling the diagnosis of retinal blood-flow abnormalities such as blood-vessel occlusion and microhemorrhages. Retinal FA requires a fluorescent dye, typically fluorescein, which is stable within blood and which can be imaged with high contrast under appropriate optical excitation. Following intra-venous injection, the fluorescent dye circulates within the blood plasma and is imaged in the eye, enabling the study of retinal blood flow. Retinal FA reveals three main phases of blood flowing within the retina: (1) the early-phase (also known as the arterial phase), where fluorescein is visible only within arteries and arterioles; (2) the intermediate-phase, where fluorescein is also apparent in the capillaries, subsequently beginning to fill venules and partially fill veins. At this stage, bright streams of fluorescent dye can be observed within partially-filled veins, corresponding to distinct laminar flow streams within blood vessels (see Fig. 1); and (3) The late-phase where all blood vessels are completely filled with fluorescent dye (see Fig. 1). Additionally, the blood vessels of the choroid, at the posterior of the retina are also highlighted with fluorescein dye, resulting in a diffuse grey background known as “the choroidal flush”.^A^ The visualization of sequential blood flow revealed by intermediate-phase FA has been used to investigate blinding diseases, such as myopia,[2] age-related macular degeneration,[3] glaucoma,[4] retinal-artery and retinal-vein occlusion,[5, 6] and carotid-artery occlusion. [7]

**Fig. 1.**
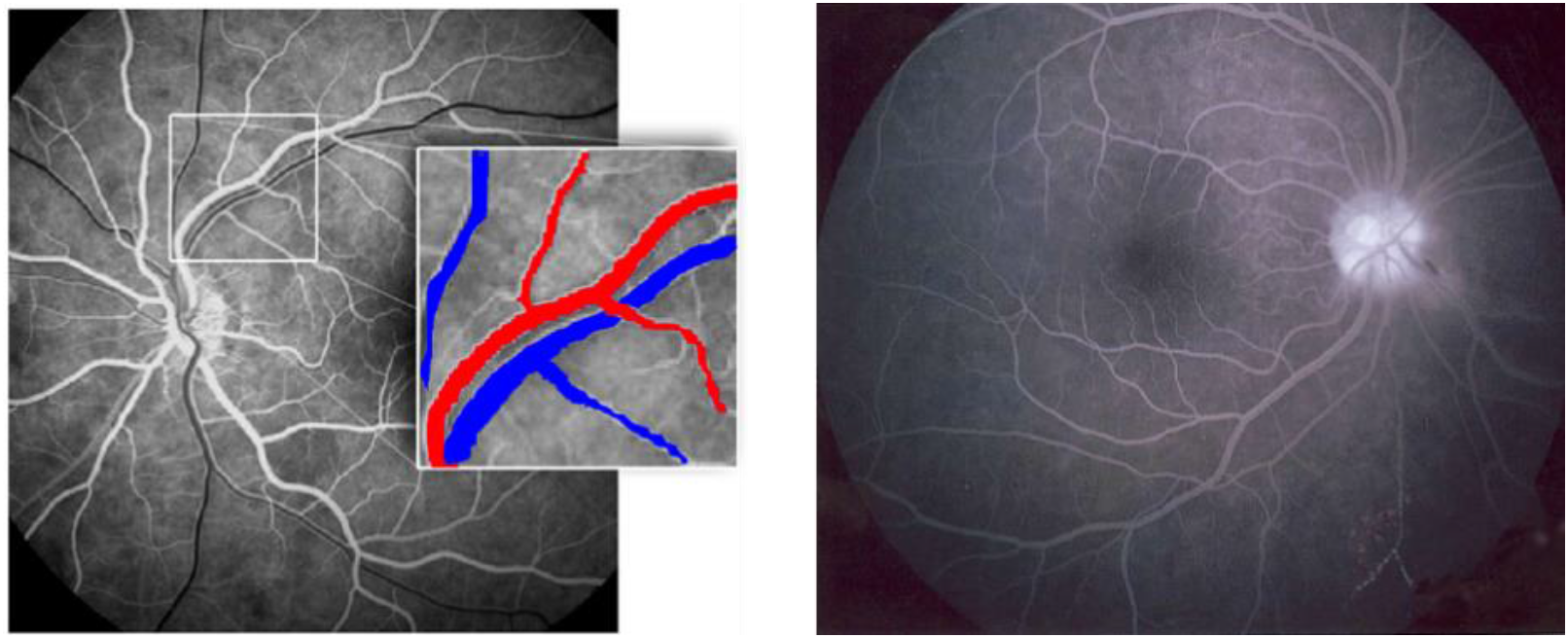
Examples of retinal FA. (Left) An intermediate-phase FA image, where arteries (labelled red in subset image) are completely filled with fluorescein whereas veins (blue in subset image) are only partially filled with fluorescein. The long fluorescein streaks within veins are indictive of multiple laminar blood flow streams within veins where blood originates from converging venules. (Right): A late-phase FA where all vessels are filled with fluorescein. Left figure reproduced under a Creative Commons Attribution License from Serlin et. al., (2013)[22]. Right figure reproduced under a Creative Commons BY-SA 3.0 license from Bacud (2007).[23].

There are however considerable risks associated with FA. Firstly, the intra-venous injection of fluorescein dye is associated with risks of severe nausea (7% of patients), skin irritation (1% of patients), and death (1 in 220,000 patients).[8–11] Alternative FA dyes such as Indocyanine Green (ICG) have been developed and function, in part, to reduce these risks, as well as to enable FA of the normally obscured choroidal blood vessels.[12] Nevertheless, there are risks associated with any intra-venous injection, and skilled medical professionals are required for administration. Further, the dyes used in FA take several hours to be filtered from the blood by the kidneys,[13] therefore preventing repeated angiography of early and intermediate-phase features. Consequently, there is considerable interest in developing non-invasive angiography techniques that can provide assessment blood-flow information, potentially with rapid repeats without the risks associated with FA.

Several non-invasive techniques that visualize retinal blood flow are available, including: Laser Speckle Contrast Imaging,[14] Laser Doppler Flowmetry,[15] Retinal Functional Imaging,[16] and several modalities of Optical Coherence Tomography (OCT), e.g. Speckle-Variance OCT and Doppler OCT. [17–19] These techniques detect or quantify blood flow; the resulting information is then processed to provide an image of retinal blood flow that is similar to a late-phase FA (see Fig. 2 for examples). However, these techniques are fundamentally incapable of generating sequences similar to early-phase and intermediate-phase FA because they do not track the circulation of a contrast agent.

**Fig. 2.**
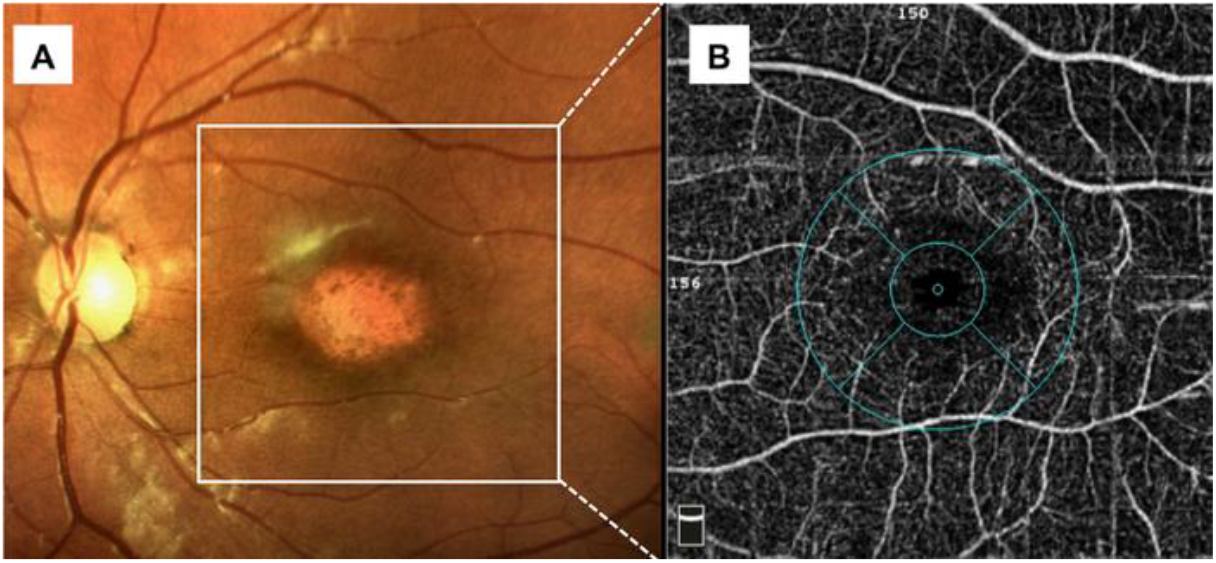
Example of OCT retinal angiography. (A) A color image of a human retina, centered on a region showing atrophy due to Stargardt disease. (B) An OCT angiography image of the highlighted region revealing a snapshot of blood flow. Note that this appears similar to a late-phase FA but does not reveal any sequential vessel filling or laminar flow. Figure reproduced under a Creative Commons Attribution License from Mastropasqua et. al., (2017).[22]

We demonstrate of a new dye-free angiography technique: Blood Oxygenation Modulation Angiography (BOMA). BOMA utilizes the oxygen-dependent absorption spectra of hemoglobin as an endogenous contrast agent, producing retinal angiography sequences with an early-phase, an intermediate-phase, and a late-phase. The oxygen-modulation contrast is created by a transient perturbation in blood oxygen saturation (OS) induced by the subject inhaling a gas mixture with reduced oxygen content.[20, 21] This OS modulation results in a temporal modulation of the absorption spectrum of circulating blood. For the retinal arteries and veins, this change is most apparent at red wavelengths (between 600 nm and 700 nm) (see Fig. 3). We imaged the change in the absorption of circulating blood using snapshot multispectral imaging, and computationally processed the recorded image sequence to yield images representing the change in OS, which can then be combined and visualized to create an angiography sequence. To the best of our knowledge, BOMA is the only non-invasive angiography technique that has the potential to visualize the intermediate-phase features of sequential blood-vessel filling and laminar flow within the retina.

**Fig. 3.**
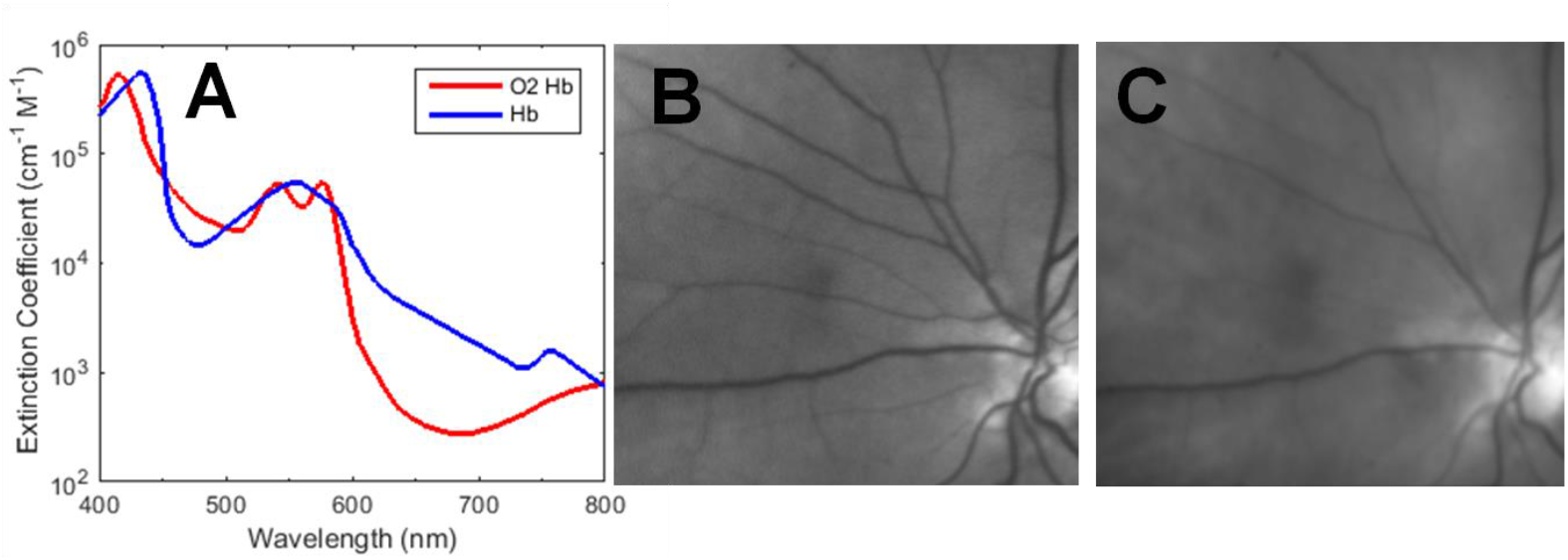
Endogenous oximetric contrast. (A) The oxygen-saturation dependent absorption spectra of haemoglobin (Hb).[24] (B) An image of retinal blood vessels at the reference 577 nm waveband (OS insensitive). (C) The same vessels imaged at the 600 nm waveband (OS sensitive), where arteries (~97 % OS) appear almost completely transparent, whilst veins (~70 % OS) absorb significantly more light and therefore appear much darker.

## 2. Principle of Blood Oxygen Modulation Angiography

For BOMA, we produce a change in the optical absorption of blood by inducing a temporary modulation of the oxygen saturation of circulating blood. This is achieved by temporarily reducing in the fraction of oxygen (FiO2) inspired into the lungs by a subject by the inhalation of a hypoxic air mixture.

Consequently, the OS of the blood within the alveolar capillaries of the lungs is decreased, initiating a transient deoxygenation wave (TDOW) of hypoxic blood, which then circulates around the body from the lungs, to other organs, including the retina. When the TDOW circulates through the retina, the absorption of blood vessels at red wavelengths (i.e. in the range of 600 – 700 nm) is increased and when the deoxygenated blood exits the retina, the absorption of blood vessels returns to pre-perturbation, i.e. normoxic, levels. This change in optical absorption due to the circulating TDOW can be visualized by measurement of the change in the intensity ratio (IR) of a retinal image at two wavebands, e.g. at a red waveband (OS-sensitive) and at a second reference waveband (OS-insensitive/isobestic). To produce angiographic sequences, the change in IR with respect to a null frame is calculated and visualized using computational image processing. In principle, the null frame selected for analysis may correspond to either normal blood-OS levels, or the peak of the TDOW in the retina (see Fig. 4). In practice, the maximum of the TDOW was found to produce the best BOMA visualizations; as discussed in Section 3.4.

**Fig. 4.**
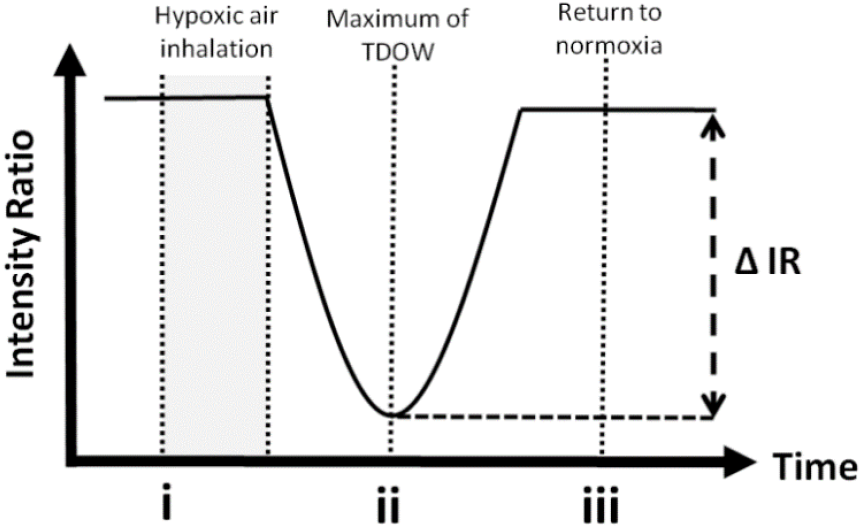
Simplified depiction of the change in intensity ratio (IR) in the retina due to a circulating TDOW. (i) Baseline normoxia (minimal absorption). (ii) Maximum of transient hypoxia (maximum absorption). (iii) Return to normoxia (minimal absorption).

## 3. Methods

### 3.1. Subject recruitment

This study was approved by the Ethics Committee of the University of Glasgow, College of Medical, Veterinary and Life Sciences. Eleven healthy human volunteers were recruited; 9 male, 2 female, age 28 ± 10 years (mean ± standard deviation). All volunteers provided informed written consent before participation and all procedures were performed in accordance with the tenets of the Declaration of Helsinki. Only subjects with no history of ocular, respiratory, or cardiovascular disease were included in the study. Retinal pigmentation is known to influence oximetry[25] and so retinal pigmentation was quantified for each subject by grading iris color on a scale of 1 (least pigmentation) to 24 (most pigmentation) according to the procedure described in Franssen et al. (2008). [26]

### 3.2 Retinal imaging with a video-rate multispectral fundus camera

Video-rate multispectral retinal image sequences were recorded using a retinal-fundus camera (TRC50 IA, *Topcon*) custom-modified with an Image Replicating Imaging Spectrometer (IRIS) and a scientific-grade CCD detector (Retiga 4000, *Qlmaging*). The IRIS is a snapshot multispectral imaging system that spectrally de-multiplexes a broadband image into eight distinct spectral wavebands onto a single detector. The eight IRIS wavebands were designed for optimal measurement of retinal blood vessel OS. [27, 28] In multispectral oximetry, snapshot imaging reduces artefacts in image quality and OS when compared to time-sequential imaging systems.[29] The IRIS has previously been applied to snapshot oximetry of retinal vessels,[30] snapshot oximetry of bulbar conjunctival and episcleral vessels,[31] and video-rate snapshot oximetry of individual red blood cells. [32] For best performance in conventional oximetry applications, the IRIS should be utilized in conjunction with a tiled spectral ‘clean-up’ filter plate to improve spectral purity, [28, 32] but for BOMA, maximal light throughput is of greatest importance so as to maximize image-acquisition rate; therefore the clean-up filter plate was not used for this study. Two IRIS wavebands were selected for image processing and analysis: 600 nm (OS-sensitive) and 577 nm (OS-insensitive, high blood vessel contrast).

Retinal illumination was provided by the fundus camera inspection lamp, operating at maximum intensity. A 530 – 610 nm bandpass filter was placed in the illumination path to constrain the spectral range of the illumination light to a single free-spectral range of the IRIS. A rotatable linear polarizer in the illumination path enabled orthogonal-polarization imaging, which greatly reduced specular reflections from blood vessels and tissue.[33] In this configuration, multispectral retinal images could be acquired at a rate of 3 Hz providing a modest temporal over-sampling to minimize image loss caused by eyeblinks, corneal glare, or motion blur from bulk eye motion. The peak optical power of the illumination configuration was measured using a paired photodiode sensor and power meter (S120C and PM120, Thorlabs). From this power measurement, the total retinal irradiance was estimated to be 1.5 mW/cm^2^. When illumination was applied continuously for 180 seconds, this corresponds to ~1/500^th^ of the photo-thermal damage threshold, and ~1/10^th^ of the photo-chemical damage threshold for the retina, as set by the International Commission on Non-ionizing Radiation Protection. [34, 35]. It would also be possible to record images with greater light intensity using flash illumination, but in this case the frame rate would then be limited by the charge rate of the flash power supply to about 1Hz providing less redundancy to images degraded by eye blinks and corneal glare. The fundus camera field-of-view was set to the maximum value of 50°. All image acquisition was controlled by *MicroManager* (University of California, San Francisco)[36] and all images were saved in 12-bit uncompressed TIFF format.

### 3.3 Experimental protocol

The left eye of each subject was selected for imaging. ^B^Approximately 20 minutes prior to the imaging procedure, the pupil was dilated by topical application of 1% W/V Tropicamide single-used eye drops (Bausch & Lomb UK Ltd.). Throughout the procedure, systemic arterial OS of subjects was monitored using a fingertip pulse oximeter (Biox 3740, *Ohmeda*), placed on the left index finger tip of subjects, operating with a data-integration boxcar-window of three seconds. Acquisition of earlobe pulse oximetry measurements using a pulse oximeter with earlobe attachment (AUTOCORR, Smiths Medical) was also attempted since this was expected to be better synchronized with the time the arrival of the TDOW in the eye, but we found that earlobe pulse oximetry data was more erratic and could not be reliably obtained. This is similar to experiences reported by other researchers attempting to use earlobe pulse oximetry.[37, 38] Multispectral retinal video acquisition and pulse oximeter data were synchronized by recording both the pulse oximeter output and ambient room audio using a microphone-equipped SLR camera (D90, Nikkon) operating in video mode. Audio cues by the fundus camera operator throughout were used as time-markers.

Subjects positioned their head on the fundus camera headrest and fixated their gaze on a red LED fixation target situated approximately thirty degrees to the right of the subject’s normal line of sight. This ensured the optic disc of the subject’s left eye was close to the center of acquired images and enabled repeated imaging of the same retinal region. Multispectral imaging of the retina was then initiated while subjects inhaled normal room air (21% FiO2, partial pressure of oxygen [pO2] ~160 mmHg) to provide a normoxia baseline measurements. After 30 seconds of normoxic breathing, subjects exhaled steadily for five seconds, and then orally inhaled a single deep breath of a hypoxic air mixture over a period of approximately three seconds. The hypoxic air mixture was a medical-grade mix of 5% oxygen and 95% nitrogen [BOC Industrial Gases Ltd] administered via a Douglas-bag gas reservoir and a flexible tube fitted with a one-way valve. Subjects then held their breath for fifteen seconds, after which subjects resumed inhaling normal room air. The pO2 of oxygen in the inhaled hypoxic air mixture was ~40 mmHg, roughly one quarter that of normoxic air, and comparable to the typical partial pressure of oxygen in venous blood. Hence, we believe that blood circulating through the alveoli during the breath hold is not subjected to the usual degree of reoxygenation that occurs during normoxia and consequently there is a transient deoxygenation of blood leaving the alveolar capillaries in the lungs. We measured a typical transient deoxygenation decrease of 5-15% using the fingertip pulse oximeter. The magnitude of the OS reduction correlated with the duration and depth of breath by the subject, but we found that with the above protocol, subjects could reliably produce an OS modulation of 5% or greater. The minimum in fingertip arterial OS was observed approximately 45 seconds after initial inhalation of the hypoxic air mixture whilst, the peak of hypoxia in the retina was recorded typically at around 10 seconds after inhalation. This difference in measurement time is commensurate with the much more direct passage of blood from the lungs to the retina, via the carotid artery, than via lungs to the fingertip.[20,21,39–41]. For those measurements we were able to make using the earlobe oximeter, the delay was typically 15 seconds (this includes both the propagation delay and a delay associated with the integration time of the instrument). Multispectral imaging was continued for a further minute after the fingertip pulse oximeter indicated a return to arterial normoxia. The typical total imaging time required for acquisition of a single dataset was approximately 180 seconds. This procedure was repeated multiple times to produce a maximum of five datasets per subject, with each dataset exhibiting a decrease in fingertip arterial OS of at least 5%. Care was taken to minimize head and eye motion of the subject during recording so as to minimize motion artefacts and glare in retinal images.

### 3.4 Image processing

For a dataset to be included for analysis, fingertip arterial OS of at least 5% was required and most images within a dataset had to be free of corneal glare, eye blinks, and motion blur. The number of datasets included for each subject are shown in Table 1.

**Table 1.** Subject demographics, valid data sets included, and results produced.

| Parameter | Subject |  |  |  |  |  |  |  |  |  |  |
| --- | --- | --- | --- | --- | --- | --- | --- | --- | --- | --- | --- |
|  | A | B | C | D | E | F | G | H | I | J | K |
| Sex [Male (M); Female (F)] | M | F | M | F | M | M | M | M | M | M | M |
| Age (years) | 34 | 21 | 54 | 25 | 24 | 23 | 25 | 24 | 20 | 26 | 27 |
| Iris pigmentation [1 = least; 24 = most] | 22 | 22 | 6 | 10 | 4 | 24 | 11 | 20 | 1 | 21 | 10 |
| Peak arterial OS decrease observed (%) | 5 | 21 | 9 | 13 | 10 | 19 | 15 | 7 | 10 | 12 | - |
| Datasets suitable for inclusion | 3 | 5 | 3 | 3 | 3 | 4 | 3 | 3 | 4 | 1 | - |
| # of TDOWs observed in retina | 3 | 5 | 3 | 3 | 1 | 4 | 3 | 3 | 1 | - | - |
| # of partially complete BOMA sequences.<sup>a</sup> | 1 | - | - | - | 1 | - | 1 | 3 | 1 | - | - |
| # of complete BOMA sequences.<sup>b</sup> | 1 | 4 | 1 | 1 | - | - | - | - | - | - | - |

<sup>a</sup> A BOMA sequence that produced angiography features, but where glare or motion artefacts did not allow for all angiography phases to be observed.
<sup>b</sup> A BOMA sequence where the early-phase, the intermediate-phase, and the late-phase were observed.

All image processing and analysis was implemented using custom algorithms implemented in *Matlab (Matlab* 2014b; *Mathworks*). A multispectral data cube was created from raw image sequences by cropping each IRIS waveband image. Each narrowband image was co-registered using a cross-correlation and de-warping algorithm to correct for minor differential optical distortions between each waveband image.[42] To compensate for eye motion within the multispectral video sequence, each image frame was co-registered to an arbitrary reference frame using a feature-matching algorithm based upon the VL_FEAT Matlab Toolbox. [43] In brief, the algorithm detects registration control-points in the reference image frame and matches them to control-points in the target image frame and co-registers all images using an affine transform.[42] Control-point features in retinal images were typically high-contrast blood vessel intersections, blood vessel perimeters, and the boundary of the optic disc. The 577 nm waveband images were used as the reference image inter-frame coregistration since this waveband exhibited the highest contrast between retinal features. Occasional corneal glare at the edge of images was removed by simply cropping all images within the co-registered image sequence.

The intensity ratio for each pixel (*x, y*) for an arbitrary frame, *n*, is:

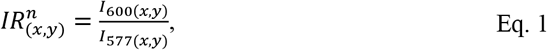

where: *I*_600(*x,y*)_ and *I*_577(*x,y*)_ are the pixel-by-pixel intensities of the 600-nm and 577-nm images. The change in intensity ratio from an arbitrary reference frame, 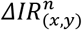, is then:

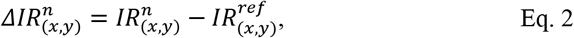

where 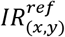 is the intensity ratio of a chosen reference frame. To reduce sensitivity to noise, the moving average of both 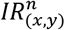 and 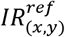 was calculated. To reduce sensitivity to noise within individual frames, a moving average boxcar of frames was applied; typically, a moving average window of 5 frames (corresponding to 1.5 seconds of data) was used. The final BOMA video sequence was generated by normalizing 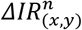 with respect to the maximum 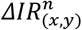 of blood vessels (not the optic disk) within the sequence. This analysis produced BOMA sequences similar in appearance to conventional FA, i.e. of bright blood vessels on a darker background.

In principle, the reference frame, 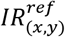 could be selected from one of several points in the sequence. For example, it would be possible to generate a BOMA sequence by using point *i* in Fig. 4, corresponding to the normoxia baseline as a reference frame, however this would mean that the BOMA sequence would be very limited in duration because the blood oxygen modulation from point *i to ii* is transient. In practice, by using point *ii* as the reference frame, the BOMA sequence produced meaningful data persistently until the end of each dataset. Further, selecting point *ii* as the reference frame was advantageous, because the subjects breathed the hypoxic air mixture between points *i* and *ii*, which typically produced motion artefacts in retinal images.

Arteries and veins were manually identified by comparison of multispectral retinal images: all blood vessels exhibit high contrast in the 577 nm waveband image, but in the 600 nm waveband image arteries appear to be highly transparent (see Fig. 1).

## 4. Results

### 4.1. Observation of TDOWs in the retina

The baseline intensity ratio (IR) of retinal regions at normoxia was strongly correlated with degree of retinal pigmentation (correlation = −0.88) (see Fig 5). A total of 29 datasets met inclusion criteria. Of these, TDOWs were observed in the retina in 26 of the 29 datasets as a characteristic decrease in retinal IR followed by a subsequent increase in retinal IR. A similar pattern was observed in fingertip arterial OS with a delay of roughly 45 seconds (see Fig. 6 for representative examples).

**Fig 5.**
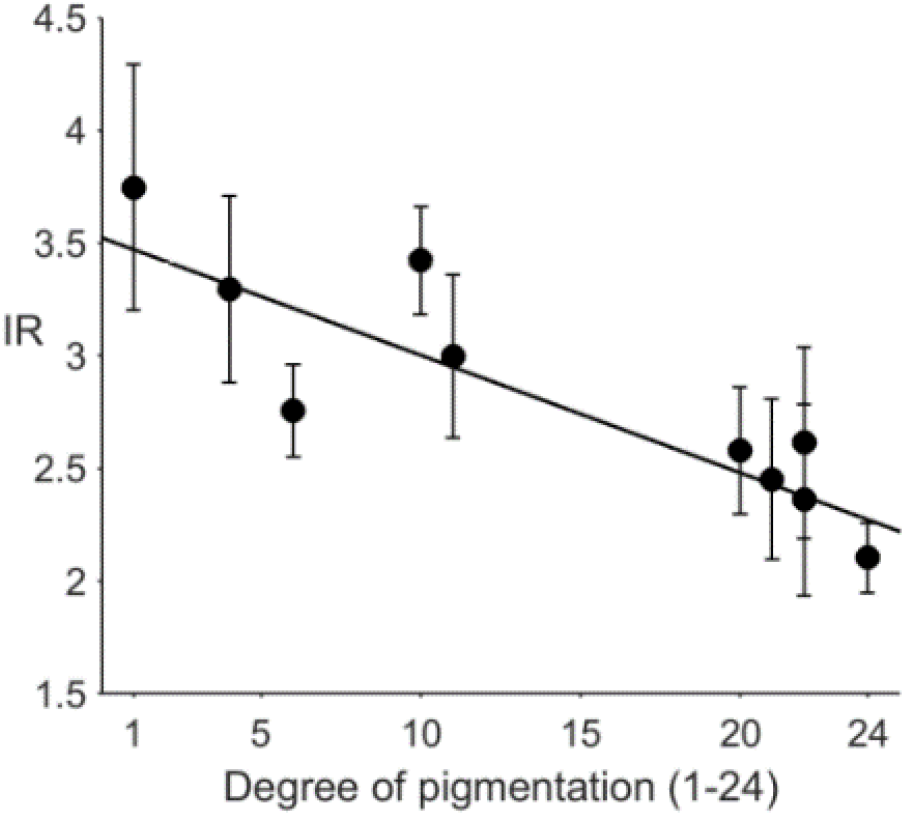
Normoxia baseline IR is inversely proportional to the degree of retinal pigmentation (correlation = – 0.88). NB: Each data point represents one subject. IR here is the average IR of retinal regions not containing any visible blood vessels or the optic disc; error bars are standard deviation.

**Fig. 6.**
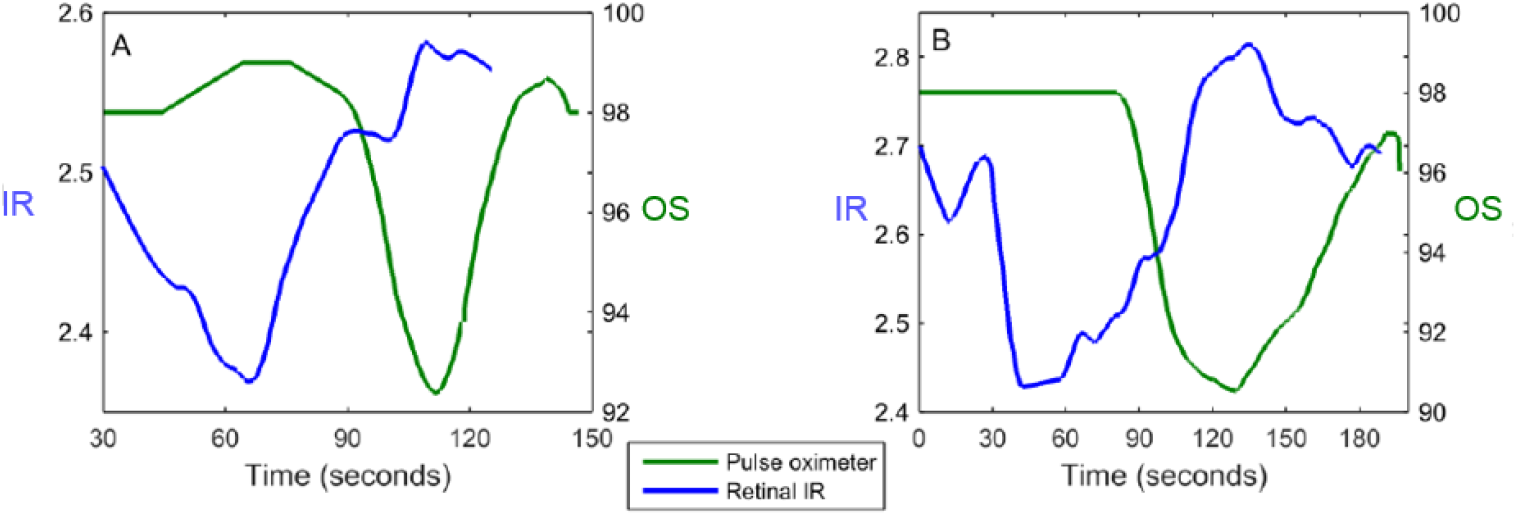
Representative characteristic TDOWs observed in retinal IR (blue line) and fingertip pulse oximeter data (green line). Each plot corresponds to a single dataset from different subjects. IR is the average intensity ratio of the entire retinal region imaged.

### 4.2 BOMA sequences and sequential vessel filling

Fig. 7 shows a comparison of a late-phase BOMA angiography image and the corresponding 577 nm waveband image of the same scene. All blood vessels are observable in the late-phase BOMA image as would be expected.

**Fig. 7.**
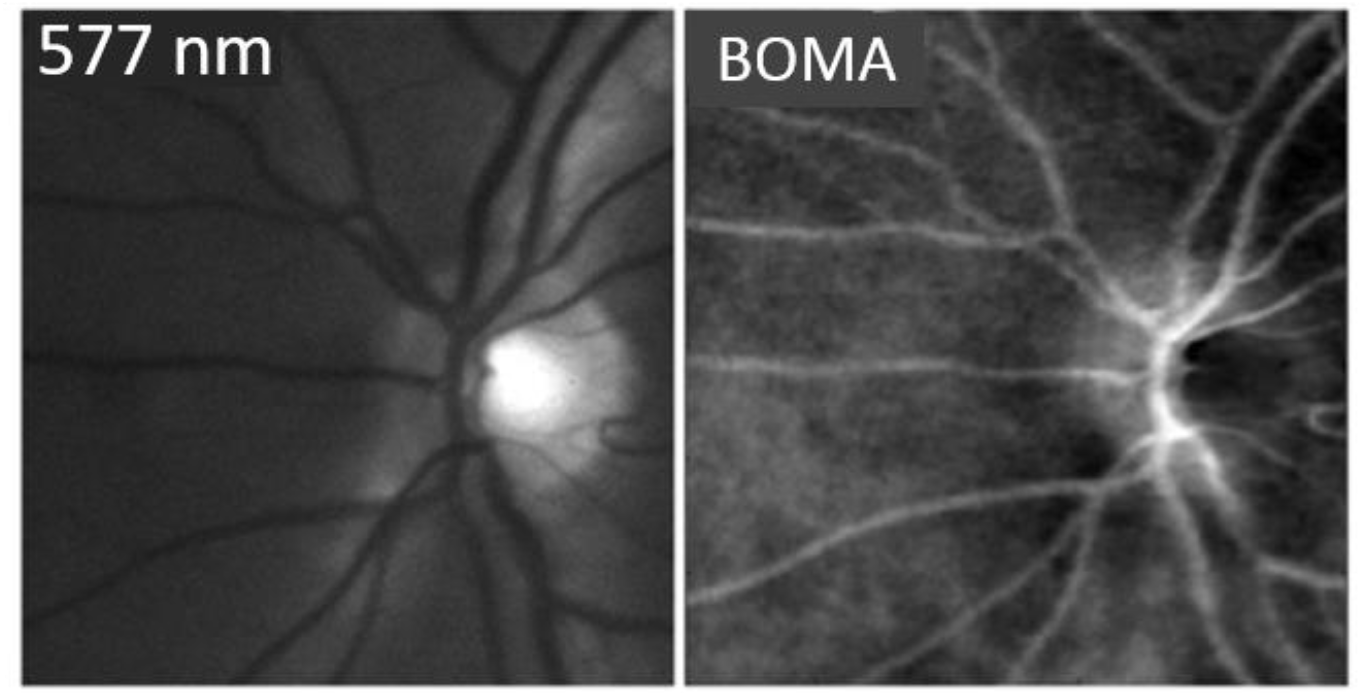
Comparison of a 577 nm waveband retinal image (left) with the corresponding late-phase BOMA angiography image (right).

BOMA sequences for three subjects are shown in Fig. 8. BOMA timepoints were selected to best display the multi-phase nature of BOMA; image-frame timestamps vary slightly because some frames were omitted from the sequence due to motion blur or eye blinks. Fig 9 shows a typical temporal evolution of Δ*IR*_(*xy*)_ for components of the eye corresponding to main arteries, main veins, and retinal regions that are dominated by background choroidal filling. Retinal arteries were observed to fill before retinal veins and the choroid, as would be expected.

**Fig. 8.**
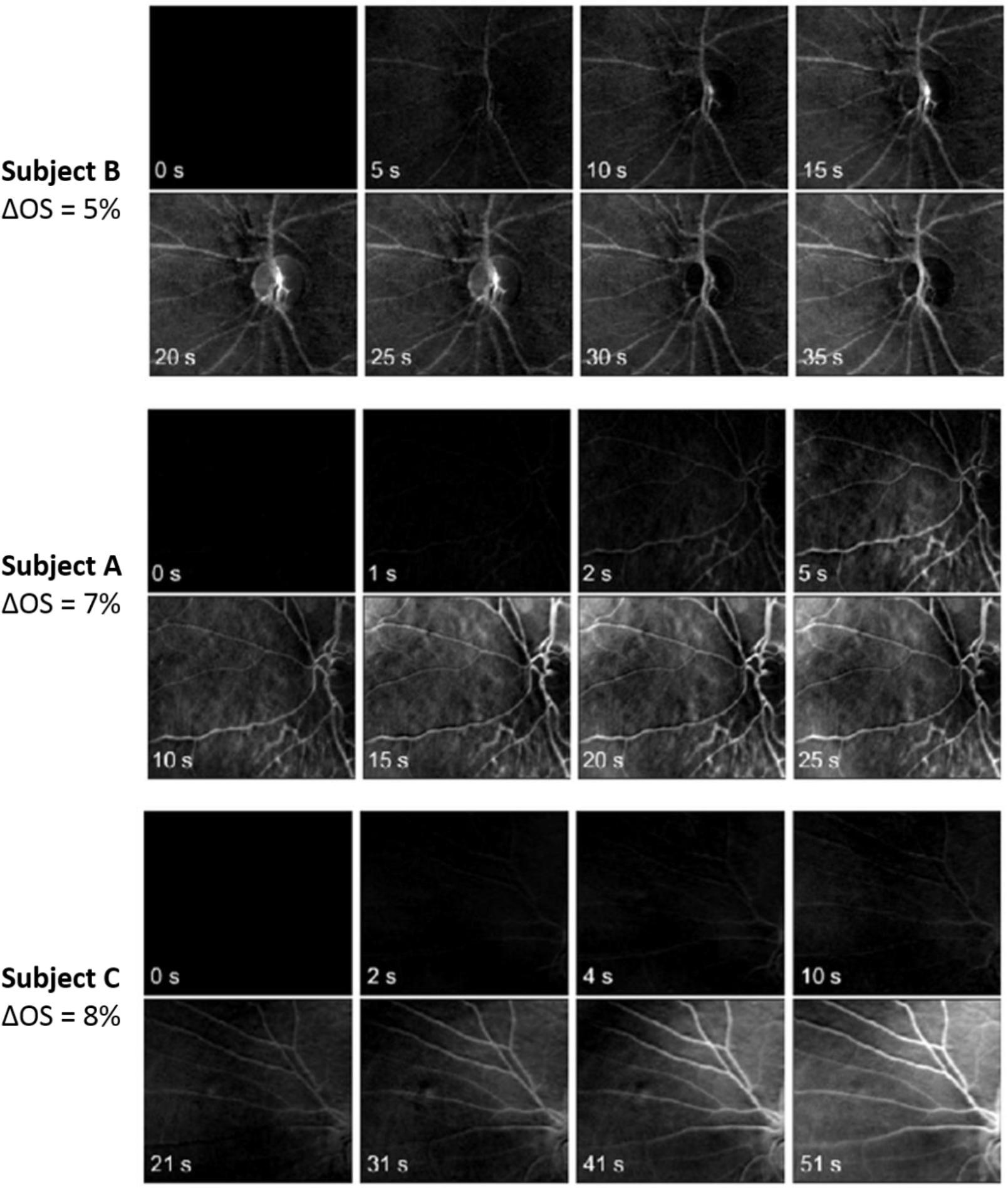
BOMA sequences for three subjects. NB: the early frames may require a high contrast digital display to observe due to weak contrast at these timepoints.

**Fig. 9.**
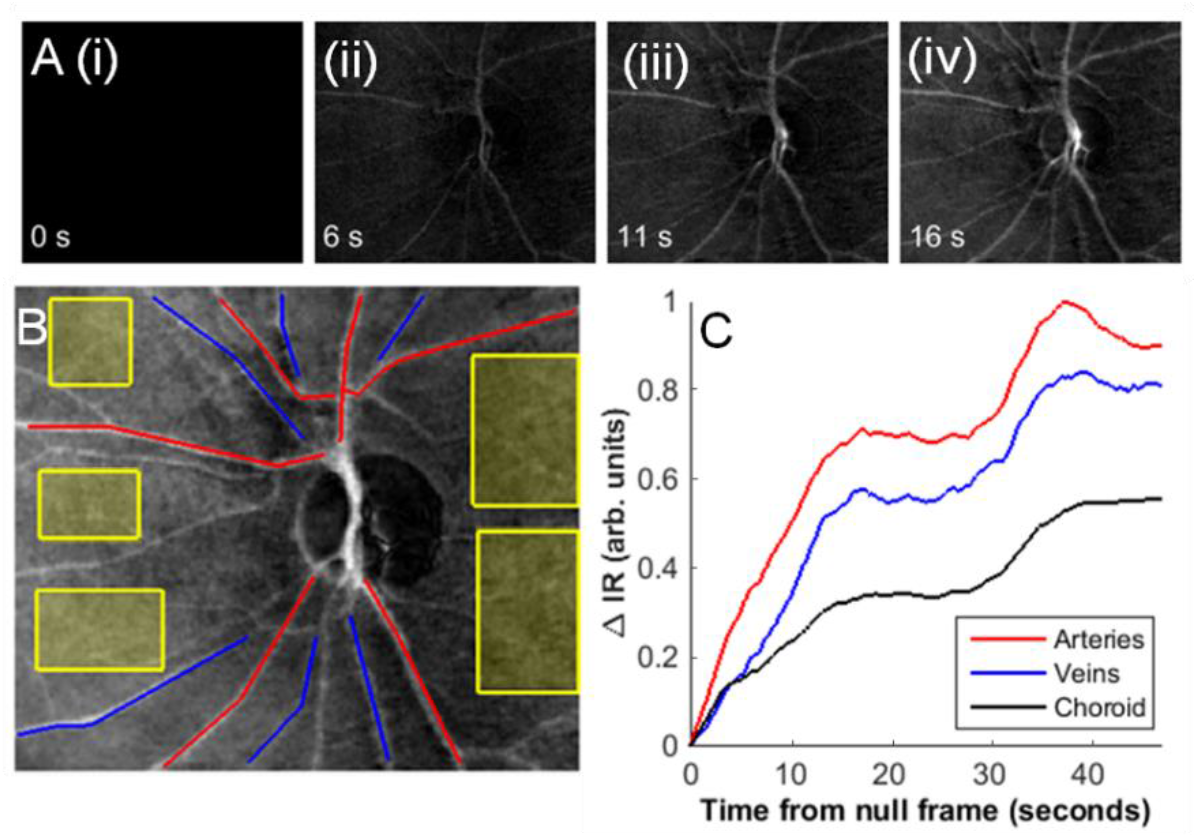
Sequential vessel filling revealed by BOMA. [A] (i-iv) Time-series of subject B showing Δ*IR*_(*x,y*)_ at (i) the reference frame, 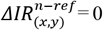; (ii) the early-phase, where arteries appear filled; (iii) the late-phase where all vessels appear filled. **[B]** The regions selected for analysis: arteries are shaded red, veins shaded blue, and choroid background regions are highlighted with yellow boxes. **[C]** Change in IR of the arteries, veins, and choroidal regions plotted vs time. Retinal arteries fill before the veins and the choroidal flush, as would be expected.

Images from four separate repeated BOMA sequences of subject B are shown in Fig 10. Good repeatability of retinal vessel angiography is clearly apparent, however there is considerable variation in the appearance of the optic disc on between and during BOMA sequences (see Fig. 11). This variable appearance of the optic disk is an artefact and is likely due to be due to small variations in eye position, which induce a relatively large 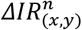 for the optic disk, compared to modest 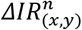 for nearby blood vessels. This susceptibility of the optic disk to artefacts may be due to the high albedo of the optic disk, it’s complex 3D morphology, and the polarization properties of retinal nerve tissue. [44–46]

**Fig. 10.**
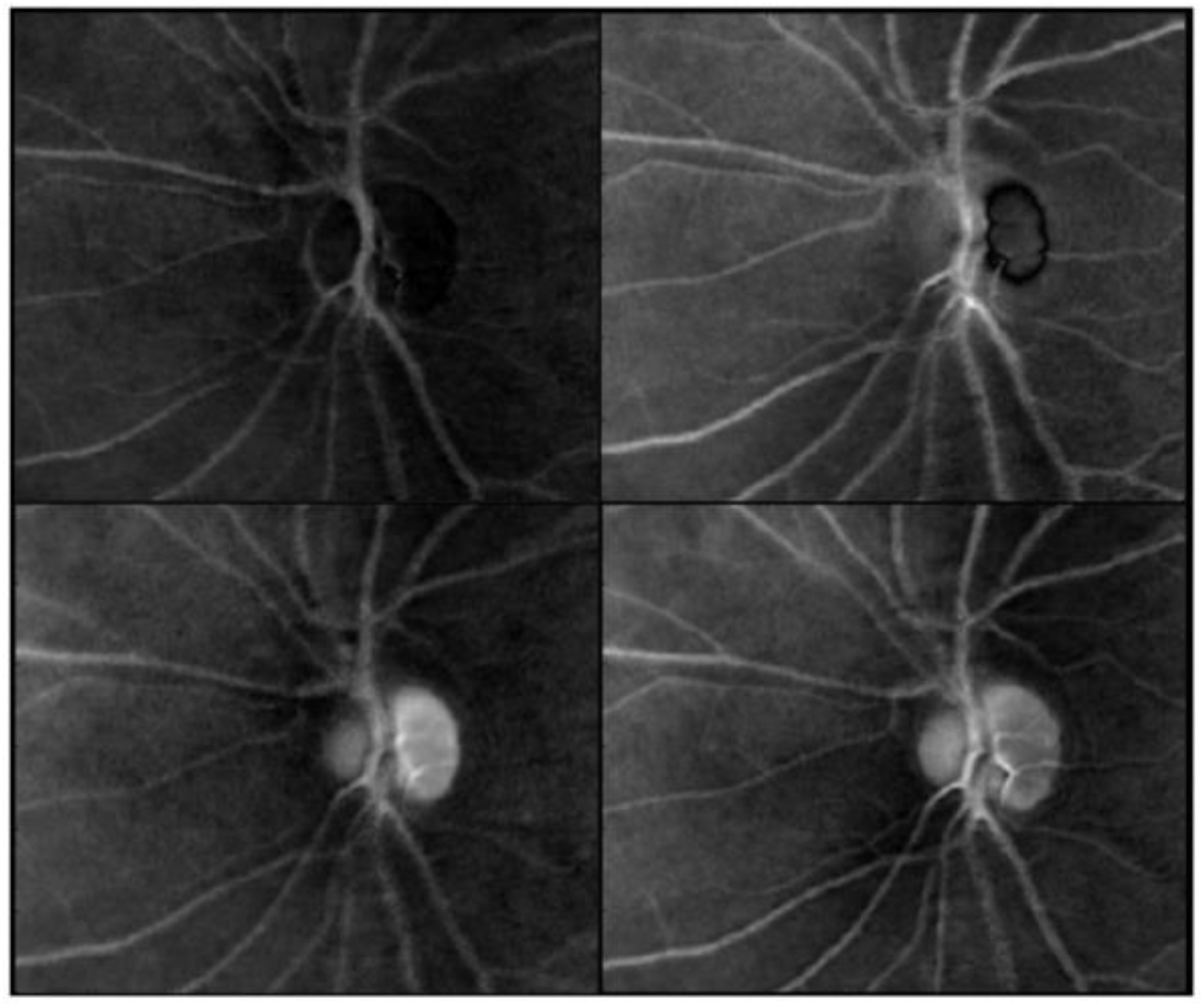
Four repeated BOMA sequences in a single subject, demonstrating that BOMA highlights blood flow in a repeatable manner. See the discussion for information on the variable appearance of the optic disk.

**Fig. 11.**
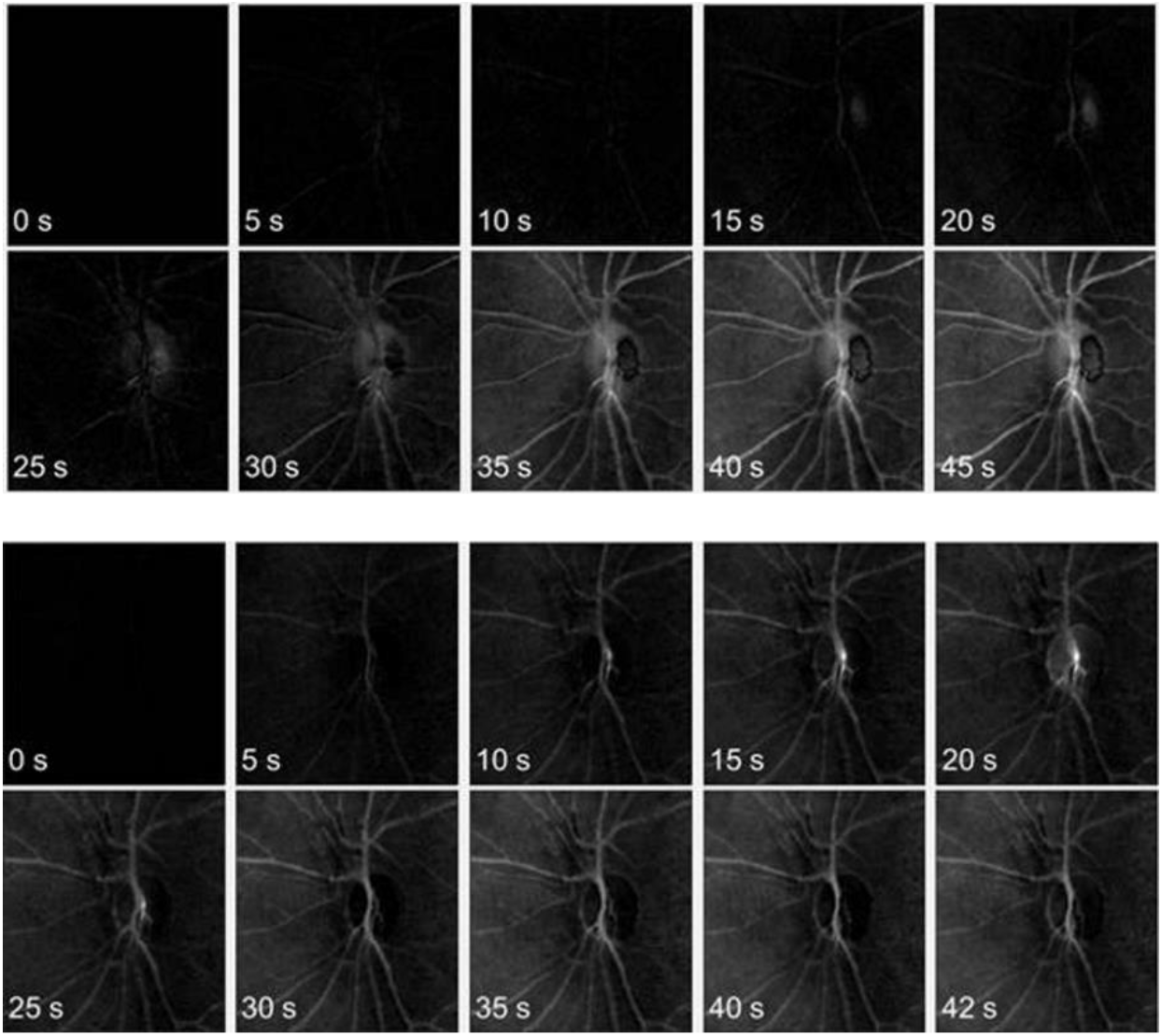
Spurious artefacts in BOMA sequences may manifest as changes in the appearance of the optic disk with a BOMA sequence. (Top) BOMA sequence where the right side of the optic disk is first spuriously bright then persistently spuriously dark. (Bottom) BOMA sequence where the optic disk is spuriously bright for a section of the sequence.

### 4.4 Potential laminar flow visualized by BOMA within retinal veins

Features consistent with our expectations of multiple laminar-flow streams within retinal veins were observed, manifesting as continuous length-wise ribbons within veins; see Fig. 12 for an example. Such laminar flow features are expected to be visualized by BOMA downstream of venular junctions where inflowing venular blood has a sufficiently high oxygen modulation contrast differential compared to the blood already present in the vein. Our expectation of this phenomena is informed by studies of laminar flow and oxygen contrast in flow channels designed to simulate multiple blood flows of different oxygenations within retinal veins (see supplementary material, Fig. S1). We do not expect to observe laminar flows within arteries since arterial blood flow diverges at branches, while in the veins, blood flow converges. In the BOMA sequence shown in Fig. 12, the fundus camera field of view was not wide enough to be able to trace the tributary venules that were the source of the laminar flow. However, the apparent laminar flow features persisted for the entire duration of the BOMA sequence (i.e. 47 seconds), which is consistent with our expectations of visualizing blood-oxygen modulation contrast. Laminar flow was not observed in the other BOMA sequences shown in this paper; possibly due to insufficient oxygen modulation contrast differentials within veins of other subjects. Further study of this phenomenon in BOMA sequences is warranted.

**Fig. 12.**
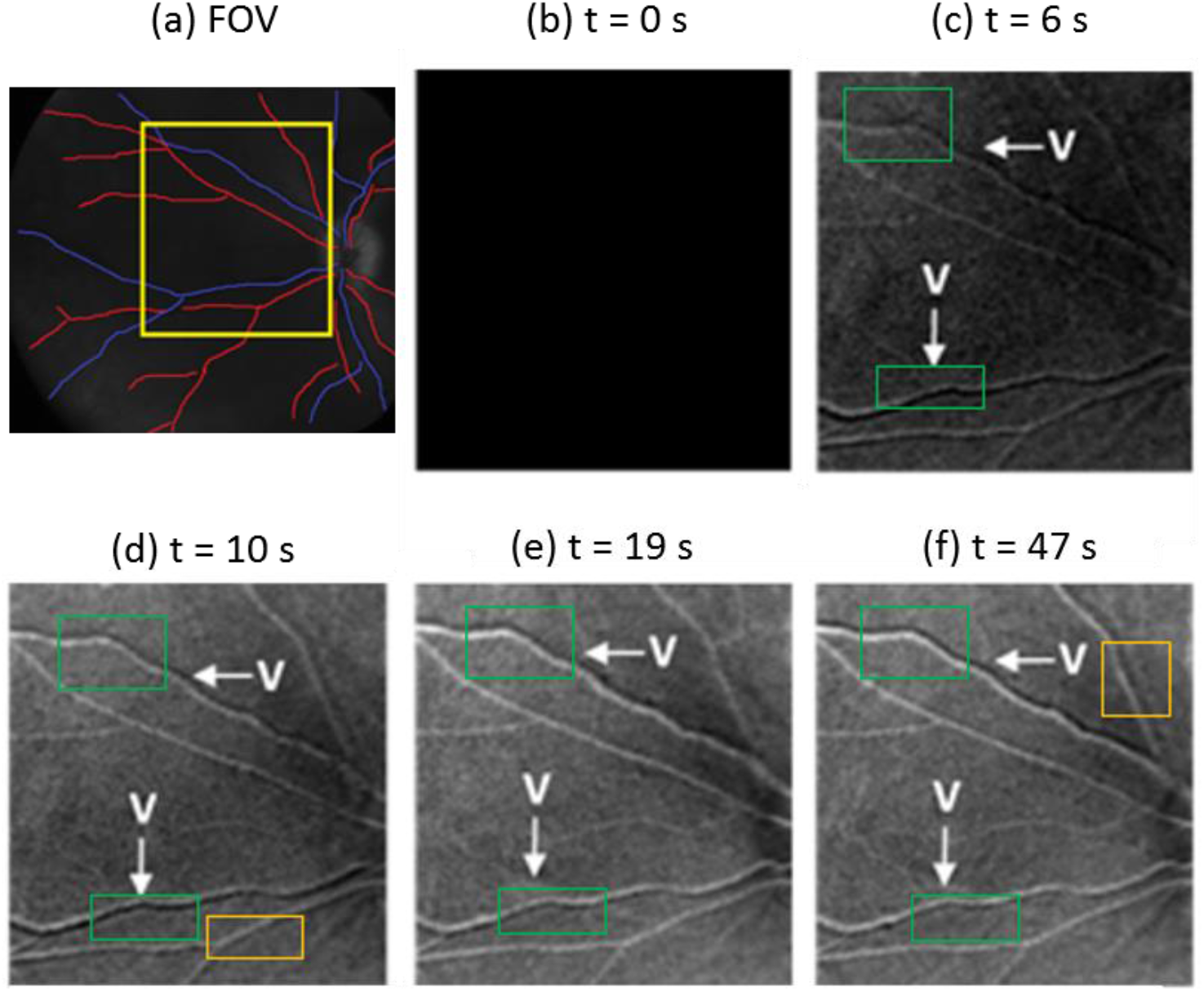
Apparent laminar flow features visualized by BOMA. (a) Retinal field of view (FOV) at 577 nm showing arteries (red), veins (blue), and the region selected for BOMA analysis (yellow box). (b – f) BOMA sequence at various time-points. Green boxes highlight regions that appear to be persistent laminar flow within veins (denoted by ‘V’). Orange boxes highlight possible transient spurious artifacts in arteries. Such arterial artefacts may arise from image misregistration errors induced by strong arterial pulsation and exacerbated by the moving average window used in image processing. N.B. early-phase arterial filling data was not available for this sequence due to corneal glare.

## 5. Discussion

Prior studies by others have used non-normoxic gas mixtures to modulate blood OS with the aim of measuring blood circulation time from lungs to other body parts, e.g. 100% N2,[20, 21] 50% CO2, 50% air mixtures,[47] <1% acetylene, 9% helium, remainder air mixtures,[40] and 100% O2.[48] We report here the first use of modulation of blood oxygenation to provide angiographic image contrast. In our study, subjects inhaled a single deep breath of a 5% O2, 95% N2 gas mixture and held their breath for 15 seconds, producing an average maximal decrease in fingertip arterial OS of 12%. This transient hypoxia is similar in magnitude to the mild hypoxia that is generally considered safe in other realms, such as the sustained ~10% decrease in OS experienced during plane travel [49–52] or up to 20% OS used in various scientific studies.[30,53,54] No adverse effects were reported by the healthy subjects from the protocol we employed here. The 30-60 seconds delay of the TDOW at the fingertip following arrival at the retina is compatible with the circulation time delays reported by other studies.[20,21,39–41]

BOMA has two primary advantages over FA: (1) it avoids the risks associated with intravenous injection of contrast agents and (2) it can be repeated on timescales of around 5 minutes opening up the possibility of repeated use.

We have observed vascular image features that are consistent with what would be expected from laminar flow of blood of differential oxygenations combining in flow channels. In supplementary Fig. S1 we show an eight-band multispectral image of flowing blood of differential oxygenation combining in a microfluidic channel *in vitro*, which shows similar laminar flow. Although there will be some diffusion of oxygen across the blood flow, from the highly oxygenated blood to the deoxygenated blood, the diffusion rate appears not to be sufficient to noticeably reduce the differential oxygenation. Similar laminar flow features were observed *in vivo* by Hendargo et al. using multispectral imaging.[29] Such laminar-flow features are not typically accounted for in studies of blood oxygenation, where analysis assumes a single uniform OS across the width of a vessel [55], although this effect could introduce some error into venular oximetry.

There are some limitations of BOMA which should be noted. Firstly, BOMA is based upon the endogenous contrast of hemoglobin, which is primarily contained within red blood cells. Therefore, BOMA is fundamentally unsuited to detection of leakage of blood plasma via microhemorrhages as commonly observed with FA.

Secondly, artefacts in BOMA could possibly arise from inter-frame blood-vessel misregistration, which would likely manifest as lengthwise vessel “shadows”, potentially similar in appearance to what we expect of laminar flow. Such misregistration errors are likely to arise from excessive bulk eye motion or from the pulsatile motion of retinal arteries. Pulsatile artefacts are however absent in veins and are unlikely to be the origin of the observed laminar-flow features in the veins. Therefore, like the spurious optic-disk artefacts, we expect the “shadowing” misregistration artefacts to be temporary in nature.[56] Future improvements to the image processing will reduce the impact of the above image artefacts.

Practical improvements can be implemented to improve data acquisition. Improving the switching between room and hypoxic air with minimize head and eye motion which often introduced glare and/or motion blur into images. Implementing high-repetition-rate flash illumination will also improve image quality by maximizing image brightness and reducing effective integration time so as to remove motion blur.

## 6. Conclusions

We have demonstrated a proof-of-concept for a new non-invasive, dye-free retinal angiography technique, Blood Oxygenation Modulation Angiography (BOMA), in healthy human subjects. BOMA exploits a temporary modulation of the OS-sensitive optical absorption of blood to act as non-invasive endogenous contrast agent. We implemented BOMA by inducing a transient deoxygenation wave (average 12% OS reduction) whilst acquiring multispectral retinal images with a snapshot multispectral imaging device connected to the output port of a retinal fundus camera. Image processing and analysis was used to generate BOMA sequences, which exhibit sequential vessel filling and laminar flow similarly to early-phase and intermediate-phase FA. Compared to other retinal angiography modalities, BOMA is advantageous because it is non-invasive, yet it can reveal flow early-phase and intermediate-phase flow features, whilst being repeatedly implemented on timescales of a few minutes. Further refinement to instrumentation, experimental protocol, and image processing will enhance BOMA image quality to be closer to that of a conventional FA.

## Supplementary Information

**Fig. S1:**
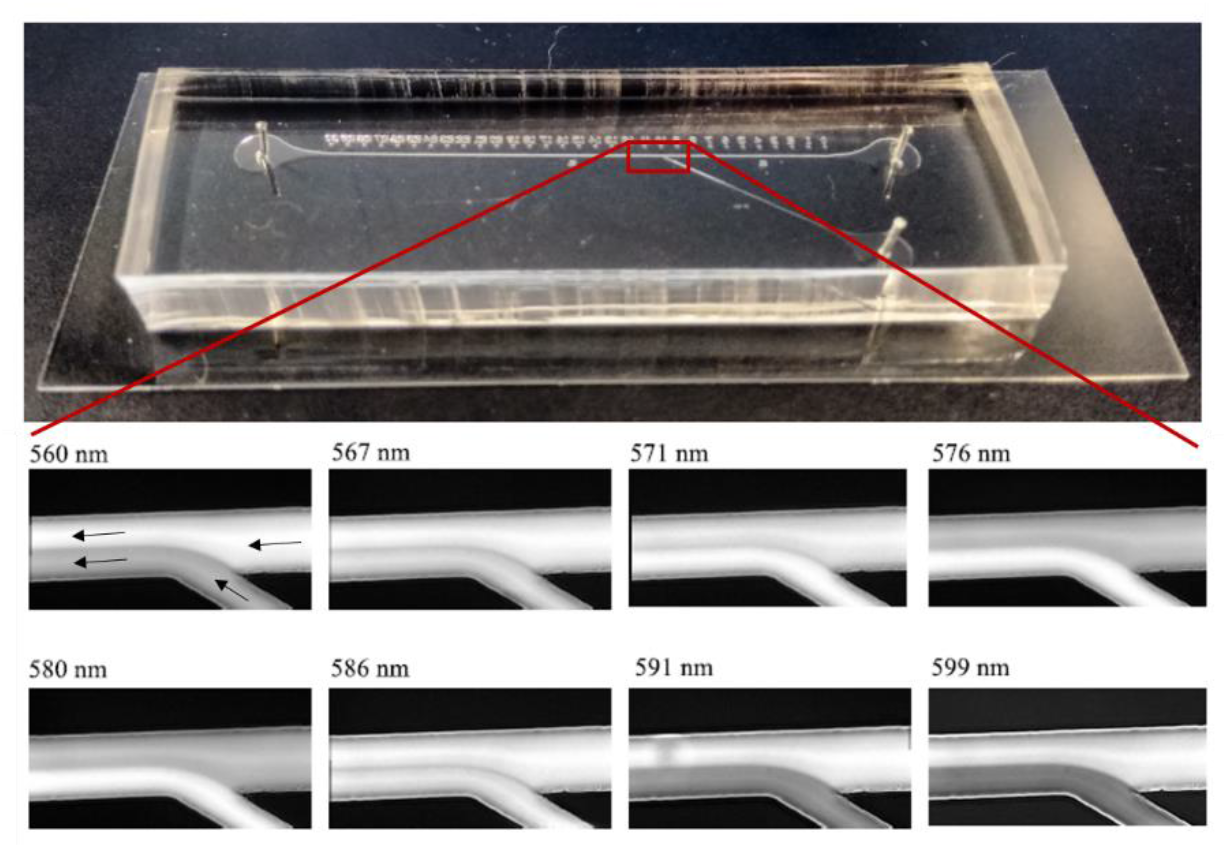
Oxygen contrast and multispectral imaging can distinguish laminar blood flows within a simulated branching vein. Top: the flow chamber, made of transparent PDMS plastic. With dimensions to mimic a branching retinal vein: the main channel 100 μm wide and the branching side-channel is 50 μm wide. (Bottom) Inverted-contrast multispectral transmission microscopy images of the flow chamber with 0% OS *ex vivo* horse blood in main channel and ~100% OS blood in the branching side-channel. Arrows annotations in the 560 nm band image indicate blood flow-direction. Laminar flow of blood can be clearly distinguished from the multispectral images. It should be noted that these images were acquired with the IRIS using the spectral clean-up filter plate, therefore the 576 nm waveband image shown in this supplementary figure is not isobestic. Full details of this experiment are available in the PhD thesis of Dr Javier Fernandez Ramos: *‘Snapshot multispectral oximetry using image replication and birefringent spectrometry’* (2017).

University of Glasgow. http://theses.gla.ac.uk/id/eprint/8162

## Footnotes

A These diffuse background characteristics of the “choroidal flush” arise because the light emitted from fluorescein within the choroidal vessels is multiply scattered by the retinal tissue. Further, melanin pigmentation in the retina acts to obscure the dense choroidal vasculature. However, in subjects with low levels of retinal pigmentation, the choroidal blood vessels can be directly observed.[57]

B Note: The left eye of each subject was selected for study in this instance because the geometry of the lab table interacted with the air inhalation equipment in a manner that made imaging the right eye challenging. In principle either eye could be used.

